# Social and reproductive benefits of group fission in wild geladas

**DOI:** 10.64898/2026.10.12.758649

**Authors:** Jacob A. Feder, Jacinta C. Beehner, Noah Snyder-Mackler, Amy Lu, Thore J. Bergman

**Author notes:** Indicates co-last authorship (Amy Lu, Thore J. Bergman).

## Abstract

Living in large, crowded groups can result in fitness costs due to increased competition and conflict. When dispersing from large groups isn’t possible, fissioning into two or more smaller ‘daughter’ groups may provide an important mechanism for mitigating these costs. Here, we leveraged 15 years of behavioral and demographic data to identify both (i) the social factors that predict group membership decisions and (ii) the adaptive consequences of group fission in the gelada (*Theropithecus gelada*), a grass-eating primate in which the risk of male infanticide – the largest fitness cost to females – is highest in larger groups. After fissioning, females tended to join the same daughter groups as their close kin, closely ranked females, and other strong grooming partners, both female and male. However, lactating females did not disproportionately join the likely fathers of their dependent infants, even though doing so is expected to prevent infanticide. Group fissions were not preceded by increases in female-female aggression or salient changes in activity budgets but were preceded by increases in infant mortality. In particular, infanticide was elevated prior to fission and plummeted after groups split. Taken together, these data illustrate how animals navigate competing social priorities and highlight the role of fission in ameliorating the costs of group-living.

## INTRODUCTION

The size of animal groups is expected to be calibrated to the costs and benefits of group-living (Alexander 1974). Due to heightened pressures at exceptionally small and large group sizes (e.g., increased predation risk and feeding competition), group size-fitness relationships often follow a ∩-shaped pattern in which medium-sized groups show better health and improved reproductive performance (Koenig 2000; Roberts and Cords 2013; Creel and Creel 2015; Markham et al. 2015; Papageorgiou and Farine 2020; Tinsley Johnson et al. 2021). Across social mammals, within-group competition often hampers reproductive output at larger group sizes (Armitage and Schwartz 2000; Jolly et al. 2002; VanderWaal et al. 2009). These large group deficits have even been observed in species that rely on shareable, leafy foods (Clutton-Brock et al. 1982; Borries et al. 2008; Kappeler and Fichtel 2024), which are theoretically expected to blunt feeding competition (but see Snaith and Chapman 2007). Beyond these energetic consequences, females in many large social groups face an additional cost: higher rates of infanticide, as males are often incentivized to join and take control of groups containing more adult females (Crockett and Janson 2000; Steenbeek and van Schaik 2001).

One way to escape these costs is for oversized social groups to fission, producing two or more groups of more optimal size (Lefebvre et al. 2003; Markham and Gesquiere 2017). Permanent fissions are rare but widespread, occurring in primates (Dittus 1988; Widdig et al. 2006; Van Horn et al. 2007; Feldblum et al. 2018), carnivores (Holekamp et al. 1993; Cant et al. 2001; VanderWaal et al. 2009), rodents (Armitage and Schwartz 2000; Waterman 2002; Manno et al. 2007), and elephants (Moss and Lee 2011). Across mammals, many fissions follow a generalized pattern in which groups gradually form spatially distinct subgroups and eventually split into two or more ‘daughter’ groups (Tokuda et al. 2014; Feldblum et al. 2018). Through these incremental changes in social interaction, individuals typically preserve their strongest social ties and form daughter groups with their maternal relatives and other close social partners (Widdig et al. 2006; Van Horn et al. 2007; Wakeford and Cords 2025a).

Despite these common behavioral mechanisms, however, the ecological and demographic challenges that trigger fission vary widely. In some populations, fissions occur when resource limitations divide groups into spatially discrete subgroups, eventually prompting their permanent split (Dittus 1988; Janmaat et al. 2009). When competition for food, space, or breeding positions is particularly intense, fissions can be further marked by elevated aggression, the forceful eviction of subordinates, and increased rates of injury and death (Ichino 2006; Archie et al. 2014; Thompson et al. 2016; Sandel et al. 2026). Fissions can also occur after destabilizing events, such as the death of a central individual (Moss and Lee 2011; Stredulinsky et al. 2021) or the arrival of a new alpha male (Winkler et al. 1984; Janson et al. 2012), the latter of which is often linked to infanticide (Teichroeb and Jack 2017). When fissions are induced by such demographic changes, group membership decisions can be more strategic. For instance, after the arrival of an immigrant male, a female might join the new dominant male or instead remain with the previous alpha male (i.e., the likely father of her most recent offspring) – a possible strategy for avoiding infanticide (Hohmann 1989; Nsubuga et al. 2008; Zhao et al. 2011). In this way, fissions allow individuals to both reshape their social environment and escape the costs they would otherwise continue to face at their current group size.

In recent years, the growing toolkit of animal social network analysis (Webber and Vander Wal 2019) has provided researchers with the means to capture the behavioral changes that precede and follow group fissions, allowing for an improved understanding of how individuals navigate group choice (Lerch et al. 2021; Wakeford and Cords 2025a; Feder et al. 2025b). However, despite clear expectations rooted in socioecological theory, data highlighting the functional benefits of fissions are surprisingly lacking. Few studies have linked fissions with their potential health and fitness consequences in long-lived animals (but see Hsu et al. 2017; Wakeford and Cords 2025b), either by measuring indirect proxies of the competitive environment (e.g., home range quality, body condition) or by directly assessing survival and reproductive output. This is likely due to the rarity of fissions – many studies are limited to single fission events, resulting in limited statistical power and generalizability. Thus, data across several groups and multiple years may be needed to detect the social and reproductive benefits of fission.

Geladas (*Theropithecus gelada*) are an ideal species for capturing both of these patterns. Geladas are cercopithecine primates that live in large, multilevel societies consisting of several reproductive ‘units’ composed of one dominant leader male, 0-3 follower males, 1-12 adult females, and their dependent offspring (Snyder-Mackler et al. 2012b). Gelada units combine to form larger aggregations called ‘bands’, in which recently fissioned units show stronger spatial and genetic affinities (Snyder-Mackler et al. 2014). This mirrors a broader pattern observed in other multilevel societies, in which units that are genetically connected through fission or dispersal remain close neighbors (Archie et al. 2006; Tong et al. 2015). In turn, familiar social units might even provide passive or coordinated defense against predators (Schreier and Swedell 2012; Camerlenghi et al. 2023) and bachelor males (Rubenstein and Hack 2004; Grueter and van Schaik 2009), perhaps because kinship and familiarity increase the potential for interunit cooperation (Wittemyer et al. 2005; Grueter et al. 2020).

Given their relatively narrow range of viable unit sizes, fissions appear to be more frequent in geladas than in other cercopithecine primates (e.g., toque macaques: Dittus et al. 1988; savanna baboons: Van Horn et al. 2007). However, within-group competition is unlikely to explain these frequent fissions, as geladas primarily consume superabundant grasses (Jarvey et al. 2018), and birth rates are unrelated to unit size (Tinsley Johnson et al. 2021). Instead, fissions may be most strongly linked to the intensity of sexual conflict and its associated costs. Females in large units experience more frequent leader male replacements (hereafter, “takeovers”) and – as a result – face higher rates of injury and sexually selected infanticide (Schneider-Crease et al. 2020; Tinsley Johnson et al. 2021). Nevertheless, accumulating evidence suggests that geladas occasionally engage in contests over seasonally available, energetically rich underground foods (Jarvey et al. 2023; Perlman et al. 2024), so nutritional constraints cannot yet be wholly ruled out as a potential driver of unit fission.

In geladas, fissions always occur following male takeovers, when multiple leader males divide relatively large reproductive units into two or more daughter units. Because these daughter units often fall closer to or within the optimal range of 5-7 females, at which both offspring and adult survival are maximized (Tinsley Johnson et al. 2021), fissions are expected to generate fitness benefits. Given the social structure of geladas, fissions may come with few associated costs, as daughter units continue to remain in large bands (300+ individuals) that appear to provide a low-cost buffer against predation (Dunbar and Dunbar 1975) and male intrusion (Pappano et al. 2012). Previous work using a small sample of fissions demonstrated that gelada units typically divide along lines of kinship (Snyder-Mackler et al. 2014). Thus, female geladas appear to have more social agency than female hamadryas baboons, whose unit membership outcomes are largely constrained by dominant males (Swedell et al. 2011). However, the behavioral processes that underlie these kin-biased unit membership outcomes have not yet been described or quantified.

Here, we used 15 years of longitudinal data to examine the social and reproductive outcomes of nine fission events in a wild population of geladas in the Simien Mountains National Park, Ethiopia. First, we used mediation models to assess whether kinship, dominance rank, age similarity, and grooming relationships jointly shaped unit membership decisions following fission. Second, we conducted a parallel analysis examining whether females similarly joined their closest male partners, particularly when they had dependent infants that were at risk of infanticide. Lastly, we examined the potential benefits of fission by determining (i) whether rates of aggression were elevated before and declined after fission, (ii) whether changes in females’ activity budgets indicated increases in social instability, and (iii) whether fissions corresponded with improved infant survival via reductions in infanticide risk. In doing so, this study captures how fissions allow individuals to reshape their social environment and improve their reproductive outcomes.

## METHODS

### Study site and subjects

Data for this study were collected from a wild population of geladas in the Simien Mountains National Park, Ethiopia. This population has been under continuous study by the Simien Mountains Gelada Research Project (SMGRP) since January 2006. Throughout this time, the SMGRP has collected demographic and behavioral data on several reproductive units that are habituated to human observers on foot. Here, we focused on the 15 most frequently sampled founding study units and their descendants that were formed via fission or fusion (see **Figure 1** for a study population history).

**Figure 1.**
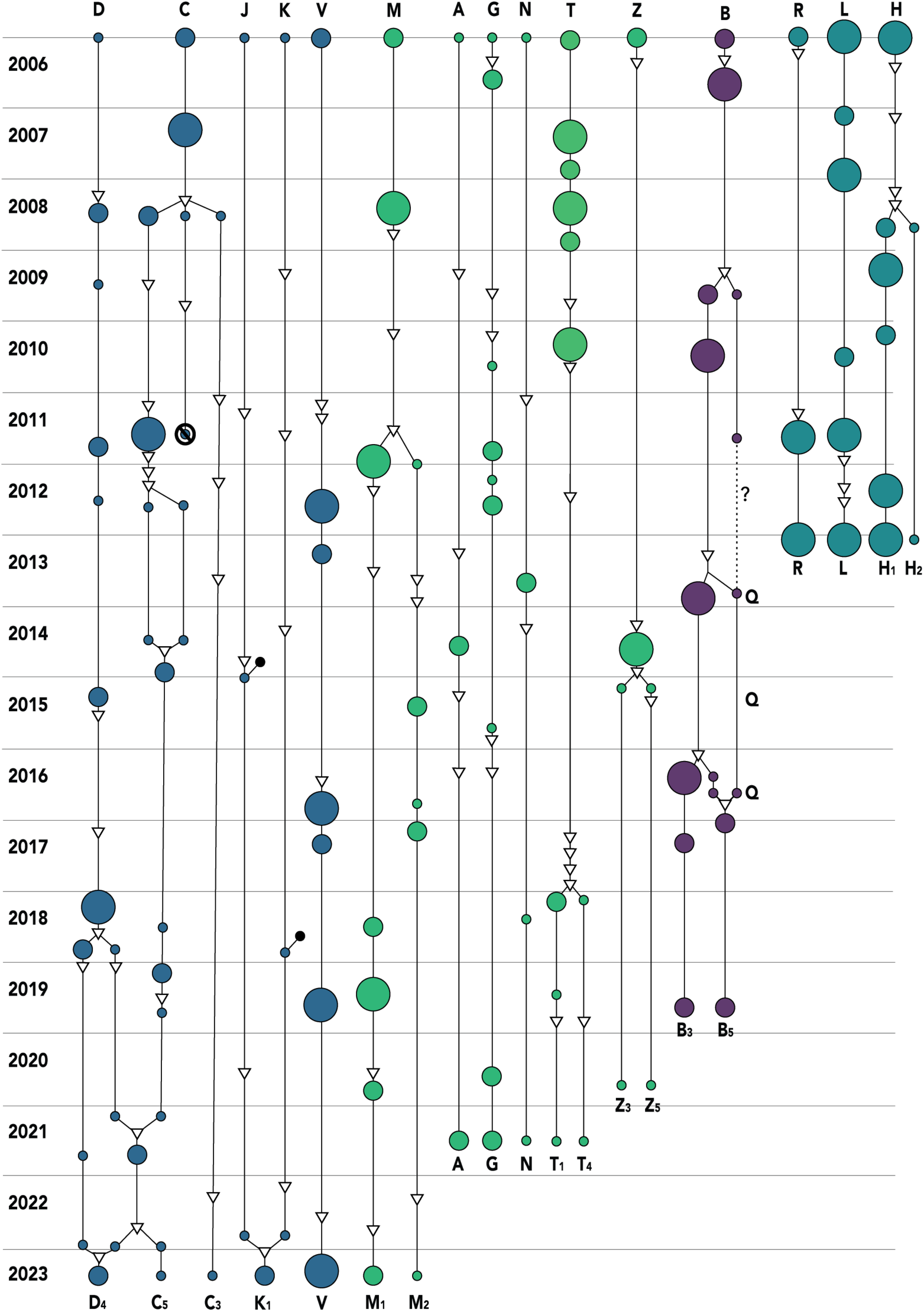
Demographic history of all study units (on the horizontal axis) across years (on the vertical axis). The size of each circle corresponds to the size category of the unit (small: 1-4 adult females; medium: 5-7 adult females; large: 8-12 adult females). Triangles indicate male takeovers, and black circles indicate instances where non-kin females immigrated into study units, which were rare (n = 2). Branching of the solid lines indicates unit fissions or fusions. Lines end when routine observation of units concluded or when units became extinct, as indicated by the ∅ symbol. The color of each unit indicates its band membership.

In total, nine units experienced fission events over the course of the study (two of which were daughter units formed during previous fissions). Of these nine fissions, two were short-lived, as the daughter units fused after two and five years of separation respectively. In another case, two daughter units briefly fused for a three-month period before splitting again into two permanent units, further highlighting the complexity of the fission process. Daughter units always remained in the same band post-fission and regularly fed in close proximity. In geladas, as in many other cercopithecine primates (Cords 2012), females are philopatric and remain in their natal units for life, except in cases of fission (le Roux et al. 2011), whereas all males disperse at 4-7 years of age (Sen et al. 2022). For these reasons, we focused primarily on unit membership decisions from the perspective of philopatric adult females.

### Behavioral data

Behavioral data were collected from 64 adult females and 32 adult males across six fission events (including the two impermanent fissions listed above). Three additional fission events predated the start of routine behavioral observations in 2009 and are thus only included in the infant survival analyses described below. Behavioral data were collected via 15-minute focal animal observations (Altmann 1974) during which grooming bouts, agonistic interactions, and behavioral states (e.g., resting, traveling) were continuously recorded. Both actor and recipient were noted for all directed social interactions. For each fission, we aggregated behavioral data across a six-year window comprising three sampling periods: 1) a stable period, which included data from 4 years before the fission to 2 years before the fission; 2) a pre-fission period, which included data from the 2 years before the fission; and 3) a post-fission period, including data from the 2 years following the fission. Altogether, these sampling periods include 2684.5 hours of focal animal observation (including 2137.1 focal hours for adult females). Notably, sampling was uneven across units (mean = 149.1 hours per sampling period; SD = 98.5 hours) and individuals (mean = 12.7 hours per sampling period; SD = 10.3).

### Social and demographic predictors of grooming

To examine the influence of social and demographic factors on dyadic grooming rates, we derived dominance ranks, relatedness estimates, and ages (known or estimated) for all females. All three of these factors have previously been implicated in shaping the strength of social bonds (Silk et al. 2006; Cords and Nikitopoulos 2015; De Moor et al. 2020) and influencing group membership decisions following fission (Widdig et al. 2006; Van Horn et al. 2007; Wakeford and Cords 2025a) in cercopithecine primates. Previous research in geladas has shown that closely related and closely ranked females form stronger social bonds (Tinsley Johnson et al. 2014). While an effect of age differences on grooming relationships has not yet been identified in this population, leader male geladas sire 80-100% of all offspring conceived during their tenures (Snyder-Mackler et al. 2012a). Thus, as suggested for other primates with marked reproductive skew, age similarity may serve as a useful heuristic when choosing grooming partners (Altmann 1979) and joining daughter groups (Van Horn et al. 2007).

Relatedness values were generated from a pedigree based on known maternal-offspring relationships and probable or confirmed paternal identities. To optimize pedigree construction, we supplemented observational and demographic data with long-term genetic records from the study population (Snyder-Mackler et al. 2014). Critically, this allowed us to infer the shared parentage of founding females in the study population, whose mothers and fathers were typically unknown. Genetic relatedness estimates (*r_w_*) were available for 66.1% of all female-female dyads (150/227) and were strongly correlated with pedigree-constructed relatedness estimates (r = 0.93).

Female dominance ranks were derived using behavioral records of agonism (including displacements, contact aggression, threat grunts, submissive behaviors) recorded during focal animal observations or during *ad libitum* daily observations. From these agonistic interactions, we calculated dominance ranks using the Elo-rating method (Albers and de Vries 2001; Neumann et al. 2011). Monthly relative ranks were then calculated using the proportion of co-resident females a given female outranked in that month, which ranged from 0 to 1. The rank difference of each dyad was then calculated as the absolute value of the females’ proportional rank difference. Although geladas show matrilineal rank inheritance (le Roux et al. 2011), rank differences are only modestly correlated with genetic relatedness in this population (Tinsley Johnson et al. 2014). Thus, including both rank differences and relatedness as predictors did not lead to issues of multicollinearity.

All adult females present at the onset of fissions (n = 64) had known or estimated birth dates from which we calculated age differences. Dates of birth were known within a week for 23 adult females born since the start of observations, while the remaining females’ birth dates were estimated using their size as immatures at the start of observation (n = 9), back-calculated from their date at maturation or first birth using population-wide age averages for these life history milestones (n = 16), or coarsely approximated from their total number of offspring (n = 16).

### Group membership analyses: female-female ties

To identify which decision rules regulate females’ post-fission group membership, we first built social network models using ‘bisonR’, which estimated the proportion of time dyads spent grooming using a Bayesian framework (Hart et al. 2023). For this analysis, we used data from the pre-fission period defined above, which should best reflect the strength of female-female relationships at the onset of fission. We treated the total time each dyad spent grooming (in minutes, undirected) as our outcome and modeled grooming relationships using a binomial distribution. By including dyadic sampling effort as an exposure variable, we estimated uncertainty in the strength of dyadic grooming relationships and propagated this uncertainty throughout our downstream analyses focused on the drivers of group choice. Specifically, credible intervals for dyads with higher sampling efforts were tighter than credible intervals for dyads with lower sampling effort, thereby alleviating noise due to variation in sampling effort. To further account for differences across sampling periods, we z-scored dyadic grooming relationship estimates within each unit.

To assess the factors that determined whether female-female relationships persisted after fission, we constructed a Bayesian structural equation model with an *a priori* structure. First, we modeled whether kinship, rank differences, and age differences impacted undirected grooming relationships among gelada females during the pre-fission period. Second, we modeled whether grooming relationships predicted whether females remained in the same unit post-fission.

Through this path model structure, we were able to (i) identify whether genetic relatedness, rank, and age predicted unit membership decisions and (ii) determine the extent to which these impacts were mediated through the strength of female-female grooming relationships (i.e., indirect effects) or independent of these prior relationships (i.e., direct effects).

Specifically, in the first path of this model, we modeled female-female grooming relationships (i.e., the log-likelihood of grooming as inferred by the ‘bisonR’ network models, standardized within each study period) using a ‘student’ distribution and included kinship, proportional rank difference, and age difference (in years) as fixed effects. In the second path of this model, we used a Bernoulli distribution to model the likelihood that a dyad stayed together post-fission (binary, yes/no) and included grooming relationship strengths, relatedness, rank difference, and age difference as fixed effects. In both components of the model, we included multimembership random effects, which are ideal for modeling undirected relationship data (Hart et al. 2022).

Here and below, we used weakly informative priors centered on zero for all fixed effects. All continuous variables were standardized prior to analysis, and posterior predictive checks indicated adequate model fits. To propagate the uncertainty in our grooming relationship measures, we ran mediation models across 20 draws from the posterior distribution of grooming relationship strengths and pooled these estimates using ‘brm_multiple’ (Bürkner 2017).

### Group membership analyses: female-male ties

We constructed a parallel structural equation model to determine whether females with dependent infants were more likely to stay with previous males than to join new males (Zhao et al. 2011). Because males often groom more with females whose offspring they (may) have sired (Moscovice et al. 2009; Städele et al. 2019), we also sought to examine whether group membership decisions were also driven by preexisting grooming relationships between females and resident males. Here, we included all dyads between adult females and males (leaders and followers) who were present at the time of fission and remained long enough after the fission to unambiguously determine their unit membership (n = 7 males who remained after fission, 53 dyads). As in the analysis above, we first examined whether the presence/absence of an infant (i.e., whether the female had an offspring under 1 year of age) shaped the strength of female-male grooming relationships. Then, we identified whether the presence of an infant and the strength of grooming relationships influenced whether females stayed with a given male. This model followed a parallel structure to the model above, in which we measured grooming relationships as a student-distributed outcome, modeled group choice as a Bernoulli outcome, included multimembership random effects, and pooled estimates across 20 draws from the posterior distributions for each grooming relationship.

### Agonism analysis

To assess the relationship between fissions and rates of aggression and submission, we constructed a Bayesian generalized linear mixed model using a Poisson distribution. Aggressive interactions included all displacements, vocal threats, and instances of physical contact, while submissive interactions included all fear barks, grimaces, as well as other behavioral indicators of submission. For this model, we treated the total count of aggressive and submissive behaviors observed during focal observations of each female as the outcome variable and included sampling period (stable, pre-, and post-fission) as a fixed effect. Given that the observed number of aggressive and submissive events is expected to increase in proportion with focal sampling effort, we included the logged number of focal hours for each female-period as an offset term. To account for individual differences in aggression given and received, we included female identity as a random intercept.

### Activity budget analysis

To assess the impact of fission on female activity budgets, we constructed a series of Bayesian generalized linear mixed models with a beta distribution. Across four separate analyses, we modeled whether the proportion of focal observation minutes that females spent (i) feeding, (ii) moving, (iii) resting, and (iv) socializing varied as a function of sampling period (stable, pre-, and post-fission). While these behavioral metrics are weak proxies of the competitive environment, changes in activity patterns might nevertheless capture nutritional constraints (Gillespie and Chapman 2001) or time management challenges (Dunbar 1992) that could trigger group fission. Because activity budgets are also expected to vary across the seasons and thus could differ due to arbitrary year-to-year differences in monthly researcher effort, we further divided behavioral observations into discrete seasons (cold-dry, cold-wet, and hot-dry seasons, as defined in Tinsley Johnson et al. 2018) and included this three-level categorical variable as an additional covariate in these models. By treating activity budgets as a binomial outcome relative to the total sampling effort for that female during that season (in minutes), we inherently weighted each observation to its corresponding observation time. As above, we included female identity as a random intercept term.

### Infant survival analysis

Demographic data for this study were derived from long-term records on births, deaths, and disappearances in this population. We included demographic data on all infants (n = 394) born during the study period (Tinsley Johnson et al. 2021) and divided them into three categories: pre-fission (infant conceived within 2.5 years before a fission, n = 51), post-fission (infant conceived within 2.5 years after a fission, n = 64), and stable (infant conceived outside of these ranges, n = 279). In doing so, we captured infants conceived within roughly one interbirth interval of the fission event (Roberts et al. 2017). We categorized infants relative to their conception dates and not their parturition dates because infants born within the first six months following a fission are presumably unrelated to the new leader male and thus vulnerable to infanticide. Moreover, pregnant female geladas often spontaneously terminate their pregnancies after takeovers, cutting off investment in offspring that would otherwise be likely to experience infanticide (Roberts et al. 2012). Using a Bayesian mixed-effects survival model in ‘rstanarm’ (Goodrich et al. 2018), we examined the impact of study period on the probability that infants survived to 1.5 years of age (the average age at weaning in this population: Lu et al. 2016).

Infants in this dataset were right-censored if they had not yet reached 1.5 years of age by the end of the study or the end of their unit’s observation.

We additionally assessed whether the likely causes of infant death varied across the same three time periods (stable, pre-fission, post-fission), focusing only on infants whose survival outcomes were known (n = 377). To do this, we constructed three Bernoulli generalized mixed models to assess three possible outcomes: 1) deaths by infanticide; 2) deaths following maternal death; and 3) deaths of unknown cause. Infants were assumed to be victims of infanticide if they died or disappeared within six months following a male takeover. While infanticide is seldom observed in the field, this period corresponds to the duration when infant mortality is most clearly elevated above baseline (Lu et al. 2021). Likewise, infant deaths were assumed to be the result of maternal loss if they coincided with the mother’s disappearance or if they occurred within three months after the mother’s death. All other infant mortality events were categorized as having unknown causes. For each model, we included whether the infant died of the respective cause and included time period (stable, pre-fission, post-fission) as a fixed effect. As above, maternal and unit ID were included as random effects.

## RESULTS

### Female-female relationships

In the first path of the unit membership analysis, we examined whether relatedness, rank differences, and age differences shaped the strength of female-female grooming relationships. As previously reported in this population, relatedness was positively associated with the strength of grooming relationships (β_kinship_ = 0.19, 89% CI = [0.07, 0.31]), and closely ranked dyads groomed each other more often than distantly ranked dyads (β_rankdiff_ =-0.24 ± 0.07, 89% CI = [-0.36,-0.11]). However, against our expectations, age difference had no discernable impact on female-female grooming relationships (β_age_ = 0.02, 89% CI = [-0.10, 0.13]).

In the second path of the unit membership analysis, closely related females were more likely to stay in the same unit post-fission (β_kinship_ = 0.36, 89% CI = [0.09, 0.64], **Figure 2a**), as were females with similar ranks (β_rankdiff_ =-0.47, 89% CI = [-0.75,-0.19], **Figure 2b**). However, age difference did not predict unit membership outcomes (β_age_ =-0.18, 89% CI = [-0.44, 0.07]). Even after accounting for the impacts of relatedness and rank similarity, female-female dyads with stronger grooming relationships tended to join the same daughter groups (β_groom.strength_ = 0.34, 89% CI = [0.04, 0.65], **Figure 2c**). The mediation model’s posterior predictions indicate, however, that the effect of grooming relationships only partially explained the total effects of kinship and rank (kinship: median percent mediated = 14.4%, 89% CI = [0.7%, 46.1%], rank: median percent mediated = 13.8%, 89% CI = [1.1%, 37.3%]; **Figure S1**).

**Figure 2.**
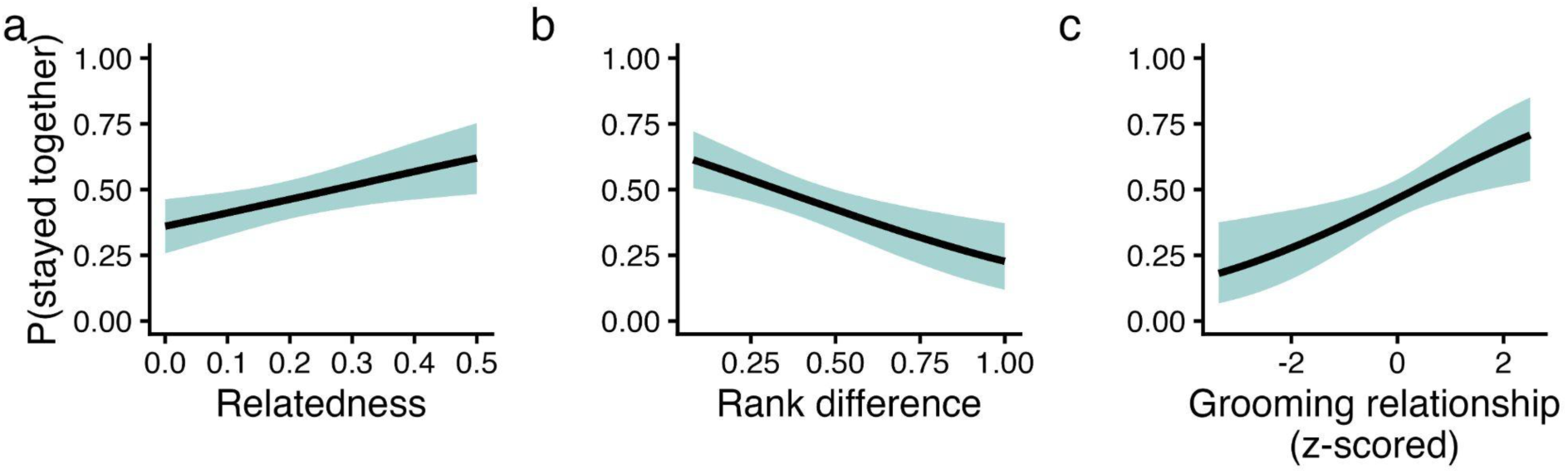
Females disproportionately joined groups containing close kin, similarly ranked individuals, and their closest grooming partners. Direct effects of (a) relatedness, (b) proportional rank difference, and (c) grooming relationship strengths on the probability that a given pair of females remained in the same unit post-fission. Ribbons indicate 89% credible intervals, while black lines indicate posterior means.

### Female-male relationships

In the first path of the mediation model, the presence of a dependent infant was unrelated to how often a given female-male dyad groomed (β_infant_ =-0.12, 89% CI = [-0.87, 0.64]).

Nevertheless, in the second path of the model, female-male dyads that groomed more often were more likely to join the same daughter group post-fission (β_groom.strength_ = 0.90, 89% CI = [0.26, 1.64]). In contrast with our predictions related to infanticide avoidance, females with young infants did not disproportionately preserve their relationships with familiar males (β_infant_ = 0.94, 89% CI = [-0.55, 2.54]). It is worth noting, however, that only 6 of 13 resident males joined a daughter unit after their respective fissions. Thus, this analysis is based on a limited sample of males, which may have precluded the detection of such an effect.

### Aggression and activity patterns

Rates of agonism were generally similar across the stable and pre-fission periods (β_pre-fission_ = - 0.12, 89% CI = [-0.26, 0.03]). However, aggression declined after a fission (β_post-fission_ =-0.47, 89% CI = [-0.64,-0.30], **Figure 3a**). Specifically, females gave or received 0.50, 0.44, and 0.29 aggressive interactions per focal-hour during the stable, pre-fission, and post-fission periods respectively. Likewise, activity budgets changed relatively little across the study periods. While females moved and rested for roughly comparable proportions of their focal observation time, time spent socializing showed incremental decreases throughout the fission process (β_pre-fission_ = - 0.12, 89% CI = [-0.16,-0.08], β_post-fission_ =-0.25, 89% CI = [-0.30,-0.21]) and time spent feeding showed concomitant increases (β_pre-fission_ = 0.08, 89% CI = [0.06, 0.11], β_post-fission_ = 0.12, 89% CI = [0.09, 0.15]). However, these changes were quite small. For instance, feeding accounted for an estimated 57.9%, 60.0%, and 60.9% of females’ activity budgets during the stable, pre-fission, and post-fission periods respectively (**Figure 3b**).

**Figure 3.**
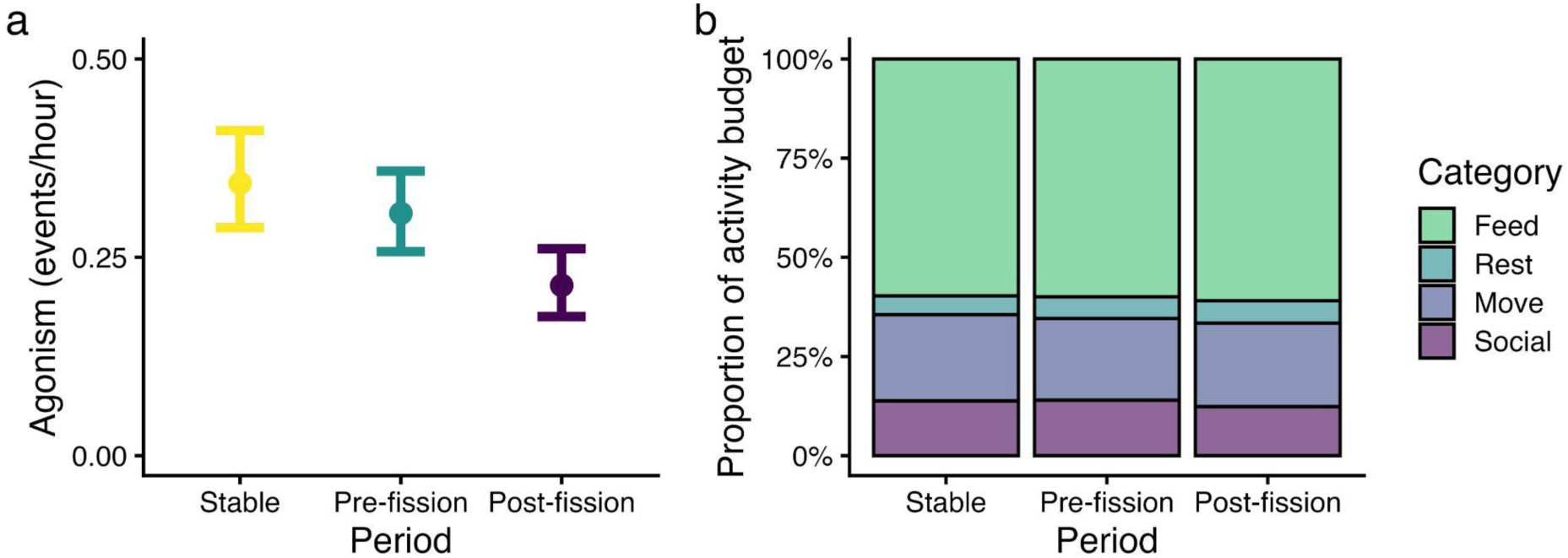
Aggression rates decreased while activity budgets remained relatively steady after fission. (a) Rates of agonism were nearly halved after units fissioned. Error bars indicate 89% credible intervals. (b) Observed proportions of time spent feeding, resting, moving, and socializing during focal observations of adult and subadult females. Although time spent socializing decreased and time spent feeding increased during the post-fission period, these changes were quite small.

### Changes in unit size

Across the entire study period (Jan 2006 - Jul 2020), fissions occurred once every 7.3 unit-years in units containing seven or more females but never occurred in units below this size. Stable gelada units (i.e., units that did not experience a fission within the following 2.5 years) contained 5.1 adult females, while units that fissioned within the subsequent 2.5 years contained 9.4 adult females on average. Thus, fissioning units are substantially larger than units drawn from the population at large (Kolmogorov-Smirnov test: D = 0.701, P = 0.0001). Post-fission daughter units, on the other hand, averaged 4.6 females, which did not differ from the distribution of stable units (Kolmogorov-Smirnov test: D = 0.135, P = 0.860). However, visual inspection of the data suggests that daughter unit sizes were often unequal, as the distribution of daughter unit sizes is approximately bimodal (**Figure 4a**).

**Figure 4.**
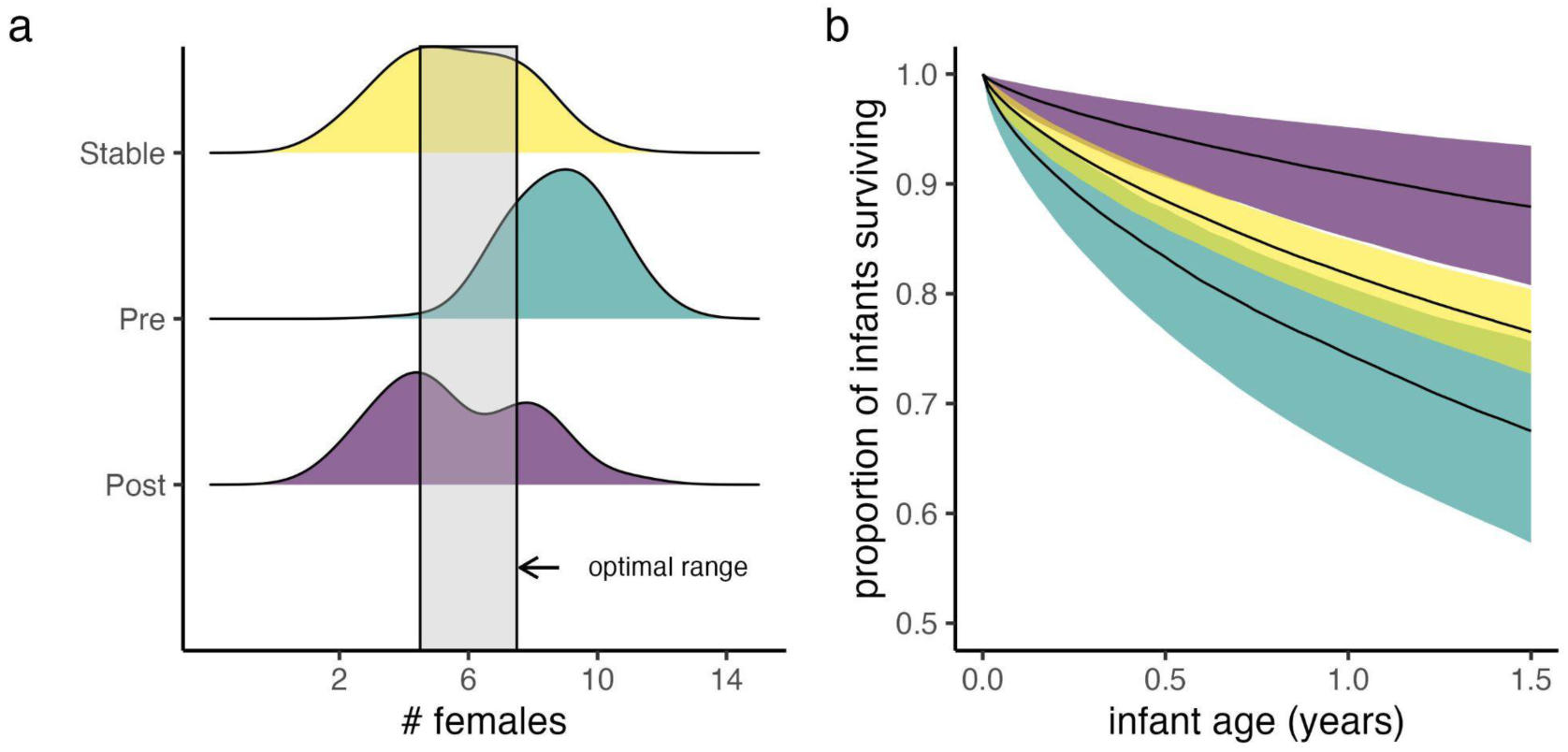
Fissions shifted reproductive units towards a more optimal size and were followed by improved infant survival. (a) Unit size distributions for females in stable, pre-fission (2.5 years prior to fission), and post-fission (2.5 years following fission) units. The gray shaded region captures the optimal unit size range, as described in Tinsley Johnson et al. (2021). (b) Survival curves of infants conceived in the 2.5 years before fission (teal), after fission (purple), or during stable periods (yellow). Ribbons indicate 89% credible intervals for survivorship to 1.5 years of age (i.e., weaning age).

### Infant mortality

Fissions were followed by reduced all-cause infant mortality: infants conceived within the 2.5 years after a fission were roughly three times more likely to survive to weaning age than infants conceived within the 2.5 years prior to fissioning (hazard ratio = 3.13, 89% CI = [1.55, 6.30]) and nearly twice as likely to survive to weaning age than infants conceived during stable periods (hazard ratio = 1.82, 89% CI = [0.98, 3.45]; **Figure 4b**). The causes of infant death similarly varied across the study periods. In particular, infants conceived following fissions were less likely to experience infanticide than infants conceived before fissions (β_post-fission_ =-3.79, 89% CI = [-6.32,-1.81]). Specifically, 23.5% of pre-fission infants were victims of infanticide, but only 1.6% of post-fission infants were. Infant deaths associated with maternal death or unknown causes, however, occurred at similar rates across all three conditions.

## DISCUSSION

Here, we documented selective group membership outcomes and improvements in reproductive success following fissions in a wild primate population. As in other female-philopatric primates, closely related and closely ranked group mates generally joined the same daughter groups, and these effects were only partially mediated through their grooming interaction patterns.

Consistent with prior research indicating that birthrates are generally unrelated to unit size in this population (Tinsley Johnson et al. 2021), fissions were seemingly not triggered by elevations in female-female conflict or increased nutritional constraints. Instead, fissions were most closely linked to infanticide, which was elevated in the pre-fission period and declined in the post-fission period. Taken together, these data suggest that fissions allow females to both preserve their strongest relationships and escape the costs of sexual conflict, thereby improving their reproductive outcomes.

### Kinship and rank predicted both grooming relationships and unit membership decisions

In general, females groomed most with their closest relatives and other closely ranked groupmates. However, age similarity did not influence grooming interactions, even though females from the same birth cohort are likely to be paternal kin. Accumulating evidence suggests that age proximity does not always underlie the strength of paternal kin bonds (Cords et al. 2018; De Moor et al. 2020), perhaps because individuals use more informative criteria when recognizing paternal kin (i.e., phenotype matching Charpentier et al. 2020). By separating paternal kin from maternal kin in future analyses, we may better detect whether and how paternal kin bonds emerge in this species.

After fissioning, females tended to form units with their close relatives and other closely ranked groupmates. Thus, female geladas may prioritize kin relationships, which have been linked to increased longevity in many cercopithecine primates (e.g., Silk et al. 2009; Brent et al. 2017). However, as in other studies of primate fissions, fissions did not always follow a nepotistic and rank-biased pattern. While most gelada units separated along lines of kinship and/or rank, one fission deviated from this general pattern. During this fission, females instead split apart along lines of age – older mothers went with one leader male, while their younger adult daughters left with the other leader male. Notably, this fission was short-lived, and the females involved reunited after roughly 2.5 years of separation. Nevertheless, this specific case highlights a key limitation of the analyses presented here, which pooled effects across all of the sampled gelada fissions and did not capture heterogeneous patterns of fission. Bottom-up, generative modeling approaches that identify which “decision rules” best match observed patterns (e.g., Lerch et al. 2021) could better capture this variability, but such an approach would ideally employ a more temporally fine-grained dataset and novel statistical approaches.

### Females did not join units containing the probable fathers of their dependent offspring

Contrary to our expectations regarding infanticide avoidance, females did not disproportionately remain with previous leader males when they had dependent offspring. Thus, female geladas do not appear to calibrate their unit membership decisions in ways that could reduce short-term infanticide risk, perhaps because the long-term benefits provided by kin bonds (e.g., Feder et al. 2025a) outweigh this risk. Likewise, gelada males did not join units containing their probable offspring but did join units containing their closest social bond partners. These patterns could indicate that both female and male geladas prioritize future reproductive opportunities over securing their past reproductive effort when selecting a new unit. However, these choices could better reflect the practical limitations on using group choice as an anti-infanticide strategy.

Because few males remained as leaders of a daughter unit, lactating females rarely had the option to escape infanticide risk altogether and join daughter units containing *only* probable or possible sire males. Overthrown males may have been similarly unable to stave off infanticide (e.g., Sen 2023), regardless of the group they chose.

Notably, the pattern we observed here contrasts with reports from many Asian colobines and gorillas, in which lactating females disproportionately form new groups with presumed sire males (Winkler et al. 1984; Hohmann 1989; Nsubuga et al. 2008). Perhaps not coincidentally, these taxa possess female-biased or bisexual dispersal patterns (Sterck 1997; Robbins et al. 2009) and/or individualistic hierarchies in which ranks are not shaped by nepotistic support (Borries et al. 1991; Robbins et al. 2005). Thus, female colobines and gorillas likely face fewer costs when severing social bonds with kin and other preferred groupmates. However, philopatry does not always preclude an anti-infanticide strategy, as blue monkey females occasionally forgo their kin relationships to join groups containing the likely fathers of their dependent offspring (Wakeford and Cords 2025a). It thus remains unclear whether these differences emerge (i) because female geladas more strongly prioritize their kin ties than blue monkeys do or (ii) because incoming leader males can more strongly constrain female group choice, as gelada units are much smaller than blue monkey groups and thus expected to be highly monopolizable (Carnes et al. 2011).

### Patterns of agonism and social activity changed after but not always before fission

Importantly, rates of agonism did not increase between the stable and pre-fission periods, suggesting that gelada fissions are not the result of elevations in female-female conflict as units grow larger. This is unsurprising, as feeding competition does not explain differences in reproductive performance across gelada units of varying size (Tinsley Johnson et al. 2021), likely because a diet focused on shareable grasses (Jarvey et al. 2018) is unlikely to generate density-dependent variation in food intake (Isbell 1991). Nevertheless, aggression rates were nearly halved following fission. Thus, even though fissions were seemingly not instigated by female-female aggression (as in Dittus 1988; Feder et al. 2025b), they may nevertheless reduce the frequency of these social conflicts. This reduction in aggression may primarily reflect post-fission decreases in unit size. Individuals residing in smaller groups often face lower rates of aggression than those in larger neighboring groups (Janson 1988), even in folivorous populations where scramble and contest competition are expected to be weak in theory but are not necessarily in practice (Blumstein et al. 1999; Koenig and Borries 2006).

By contrast, time spent feeding increased incrementally throughout the fission process, even before fissions were complete. While these changes could be a sign of increased nutritional constraints pushing larger gelada units towards lower-quality foods, we consider this explanation unlikely, as the gelada diet is highly homogenous (Jarvey et al. 2018) and these changes were quite small (1-2% of total observation time). Instead, these patterns could likewise be a mere byproduct of decreases in unit size. Both within and between populations, females in larger groups tend to spend more time engaging in grooming than females in smaller groups (Lehmann et al. 2007; but see Grueter et al. 2013). Thus, changes in unit size could have facilitated both reductions in social activity and commensurate increases in time spent feeding, as decreases in one activity must be met by increases in at least one other. These changes could have been observed even in the pre-fission period if individuals ceased investing in relationships that would later be severed and allocated this free time towards increased feeding. However, more fine-grained data on food intake or physical condition are needed to wholly rule out any energetic drivers of fission in this population.

### Fissioning improved infant survival via reductions in infanticide

Infants conceived after fissions were more likely to survive than infants conceived before fissions. By definition, many pre-fission infants experienced a takeover during their early life and thus faced an elevated risk of infanticide compared to those conceived afterwards. However, because these pre-fission deficits and post-fission improvements remained evident even when offspring conceived during stable periods were treated as the reference category, the observed differences are unlikely to be a mere artifact of study design. These improvements in infant survival were largely attributable to short-term reductions in infanticide risk. Post-fission units typically fell within or closer to the optimal unit size range (Tinsley Johnson et al. 2021), and infants conceived after fission were much less likely to experience infanticide than infants conceived during pre-fission and stable periods. By contrast, the frequency of infant deaths associated with maternal death or unknown causes remained unchanged across these sampling periods. In this way, fissioning reduced unit size and was associated with improved reproductive outcomes, as has often been presumed for other oversized primate groups (e.g., Dittus 1988) but rarely tested (Wakeford and Cords 2025b).

However, it remains unclear whether the benefits of fission are experienced equally by all daughter units. For instance, smaller daughter groups show improved reproductive output in red-tailed monkeys (Struhsaker and Leland 1988), while larger daughter groups show more pronounced gains in blue monkeys (Wakeford and Cords 2025b). Likewise, in meerkats, larger dispersing cohorts are more likely to establish breeding territories than smaller dispersing cohorts (Maag et al. 2018). In one extreme case from our study population, a too-small daughter unit went extinct within two years after its fission (**Figure 1**). Similar post-fission extinctions have been noted in chimpanzees (Feldblum et al. 2018) and in capuchins (Janson et al. 2012). In such cases, fusing with other groups could serve as a “last resort” for preventing extinction (Dittus 1987; Isbell et al. 1991; Takahata et al. 1994). While unfamiliarity with outgroup individuals and resulting collective action problems might hinder successful fusion in many social systems (Jaffe and Isbell 2010), merging with other units might be more feasible in geladas and other multilevel taxa. Gelada units residing in the same band already hold tolerant relationships and often contain relatives (Snyder-Mackler et al. 2012, 2014), and fusion has occurred relatively frequently in the gelada population (n = 5 fusions, **Figure 1**).

### Implications for the evolution and maintenance of multilevel societies

In geladas, daughter units continue to remain in the same band as highly tolerant neighbors, forming an intermediate level of social organization called a “team” (Snyder-Mackler et al. 2012b). This process generates a spatial pattern and sociogenetic structure that mirrors those observed in other multilevel taxa, such as elephants (Archie et al. 2006), killer whales (Bigg et al. 1990), zebras (Tong et al. 2015), and humans (Hamilton et al. 2007). Amicable fissions may have even been a catalyst for the evolution of many multilevel societies, enabling selective advantages for individuals that were more tolerant of their former groupmates. In particular, if kinship and shared histories among neighboring units improve coordination against extrinsic threats (Camerlenghi and Papageorgiou 2024), this could provide both direct (i.e., reduced predation and infanticide) and indirect fitness benefits (i.e., inclusive fitness gains, which could be minuscule across dyads but considerable in the aggregate).

In addition to these spatial and genetic associations (Snyder-Mackler et al. 2012b), familiar groups embedded within the same bands often retain common vocal signatures (Gero et al. 2016; Fischer et al. 2020; Painter et al. 2024) and might even share microbes despite minimal direct contact (Perofsky et al. 2017; Trosvik et al. 2018). As time passes post-fission, however, daughter units should become increasingly dissimilar in their genotypes and phenotypes, eventually regressing to the mean difference observed across the study population. Future analyses on the gelada study population will aim to capture this process of daughter unit differentiation, identify the “glue” that keeps units together despite these changes, and further examine the adaptive consequences of living in a kin-structured multilevel society.

## ETHICS

Research on geladas in the Simien Mountains National Park, Ethiopia was conducted with the permission of the Ethiopian Wildlife Conservation Authority and was approved by the IACUCs of the University of Michigan (PRO00008871), Arizona State University (20-1754R), and Stony Brook University (773805).

## DECLARATION OF AI USE

We have not used AI-assisted technologies in creating this article.

## AUTHOR CONTRIBUTIONS

Conceptualization, J.A.F., J.C.B., A.L., T.J.B.; Data Curation, J.C.B., T.J.B., N.S.-M., and A.L.; Formal Analysis, J.A.F.; Funding Acquisition, J.C.B., T.J.B., N.S.-M., and A.L.; Investigation, J.A.F., A.L., T.J.B.; Methodology, J.A.F., A.L., and T.J.B; Project Administration, J.C.B., T.J.B., N.S.-M., and A.L.; Resources, J.C.B., T.J.B., N.S.-M., and A.L.; Supervision, A.L.; Visualization, J.A.F.; Writing – Original Draft, J.A.F., A.L., and T.J.B.; Writing – Review & Editing, J.A.F., J.C.B., T.J.B., N.S.-M., and A.L.

## ACKNOWLEDGMENTS

We would like to thank the Ethiopian Wildlife Conservation Authority for permission to conduct research on geladas in the Simien Mountains National Park (SMNP), Ethiopia. We would also like to thank the wardens of the SMNP (Berhanu Gebre, Maru Biadglegn, Abebaw Azanaw Haile), the core field team members who collected these long-term data (E. Jejaw, A. Fanta, S. Girmay), and the many researchers who further supported data collection in the field (J. Jarvey, M. Gomery, L. Morris, T. Regan, P. DeLacey, P. Clark, E. Sloan, L. Babbitt, and M. Melton).

